# Achieving cell-type specific transduction with adeno-associated viral vectors in pigeons

**DOI:** 10.1101/2025.04.29.651184

**Authors:** John Tuff, Kevin Haselhuhn, Tatjana Surdin, Stefan Herlitze, Marie Ziegler, Onur Güntürkün, Noemi Rook

**Author notes:** corresponding author Dr. Noemi Rook, Institute of Cognitive Neuroscience, Faculty of Psychology Department Biopsychology, University of Bochum, Universitätstraße 150, 44801 Bochum, Germany. shared first authorship (John Tuff, Kevin Haselhuhn).

## Abstract

Birds are valuable models for studying learning, cognition, song, and vision, yet tools for controlling and recording brain activity with millisecond precision remain underutilized in avian research. Advances in methods such as chemogenetics, optogenetics, and in vivo imaging have transformed rodent studies but require gene delivery techniques, like adeno-associated viruses (AAVs), in non-transgenic species. This study validates AAV tools for precise gene expression in pigeons. We identified AAV8 as a highly effective vector, demonstrating strong neuronal tropism and anterograde/retrograde transgene expression, while AAVretro was ineffective. The CaMKIIα promoter and mDLX enhancer enabled cell-type-specific expression, targeting predominantly excitatory and inhibitory neurons, respectively. Additionally, we established proof of concept for the expression of NpHR (a chloride pump) and demonstrated the functionality of conditional gene expression systems, including Cre/loxP and Tet-On/Tet-Off. These advancements expand the genetic toolkit for pigeons, facilitating precise manipulation of neural circuits and enabling future studies on complex avian behaviors and brain functions. By bridging molecular tools and avian neuroscience, this work paves the way for comparative and translational research, offering insights into the neural basis of cognition and sensory processing in birds.

## Introduction

Avian species have emerged as valuable models in neuroscience research due to their diverse cognitive abilities and specialized behaviors. For instance, crows display cognitive skills comparable to those of primates [1–3], while zebra finches are widely used in studies of vocal learning, offering insights relevant to human language [4–6]. Pigeons, on the other hand, are renowned for their exceptional visual processing and navigational abilities [7–9]. Despite the distinct organizational differences between avian and mammalian brains [10], recent research suggests that certain principles of sensory system organization are conserved across species [11]. This growing body of evidence highlights the value of comparative studies, which offer critical insights into how brain functions emerge from neural structures in both birds and mammals [12].

Especially with the emergence of groundbreaking techniques like optogenetics, which enable precise control over neuronal activity, the possibilities for advancing neuroscientific research are rapidly expanding. Optogenetics can be used to control and manipulate neuronal activity using genetically modified light-sensitive proteins, such as ion channels (e.g. channelrhodopsin) or chloride pumps (e.g. halorhodopsin) with millisecond accuracy and cell-type specificity, which cannot be achieved with traditional manipulation methods such as pharmacology or electrical stimulation [13,14]. Through its growing popularity, optogenetics has been established in a wide variety of model species, most prominently in mammals, including rodents [15], primates [16] and ferrets [17], but also in birds, such as zebra finches [18] and pigeons [8]. In contrast to rodents, transgenic bird models are not yet available, making gene transfer methods essential for the successful application of optogenetics in avian species. A widely used approach is viral gene transfer, which employs vectors such as adeno-associated viruses (AAVs) or lentiviruses. These vectors can be administered locally or systemically [19], delivering genetic material encoding the selected opsin. Once inside the target cells, the genetic material is expressed, allowing the opsin to integrate into the cell membrane. When selecting a viral vector, it is essential to consider several factors, including the type of virus, the appropriate serotype, and the choice of promoter system [19]. In the context of optogenetics, adeno-associated viruses (AAVs) are commonly used, often with AAV2 genomes pseudotyped with various AAV capsids to enhance targeting specificity. The efficiency of viral vectors is highly dependent on the viral serotype, which can vary significantly across different tissues and species [20,21]. For instance, a recent study in pigeons compared the transduction efficiencies of AAV1, AAV5, and AAV9. The results showed that AAV1 reliably achieved robust transgene expression, AAV5 produced virtually no expression, and AAV9 had neurotoxic effects [8]. Another key consideration in selecting an appropriate viral vector is choosing the right promoter system to target specific neuronal populations [19]. Promoters are included in the gene sequence within the viral vector and are vital for opsin expression. Different promoters are for example able to drive opsin expression in either all cell types (e.g. CAG) or specifically in certain cell types (e.g. CaMKIIα for excitatory cells or mDLX for GABAergic cells) [16,22]. In pigeons, behavioral manipulations to date have exclusively utilized the widely used synthetic CAG promoter, which is driving robust and ubiquitous gene expression in a variety of cell types and organisms [8]. To advance future studies, it is essential to demonstrate that other promoter systems can reliably drive gene expression with comparable specificity to that observed in mammals. These complexities highlight the need for dedicated investigations in species where the properties of viral vectors remain less well understood. In this study, we evaluate the efficiency of various viral vectors in transducing different regions of the pigeon brain. Building on previous findings that identified AAV1 as the most effective construct in pigeons, we compared its performance to AAV8, a serotype known for reliable expression in other species. Furthermore, we examined the efficacy of several promoter systems, including CaMKIIα for targeting CaMKIIα-positive cells, mDLX for GABAergic cells, as well as the Cre-loxP and Tet-off systems for conditional expression.

## Results

### Assessment of AAV8 for Gene Transfer in Pigeon Brains

Previous studies have highlighted significant differences in the efficiency of adeno-associated virus (AAV) serotypes in transducing cells in avian brains [8,23]. For example, Rook et al. (2021) found that AAV1 is highly effective, while AAV5 and AAV9 result in little to no expression [8]. To expand the existing toolkit, we decided to investigate the suitability of AAV8, as previous studies in rodents reported a transduction preference for non-neuronal cells [24,25]. We injected AAV8 into various brain regions and observed widespread expression in areas such as the hippocampus (Fig. 1a, b), septum (Fig. 1a, c), mesopallium (Fig. 1d, e), nidopallium (Fig. 1d, f), entopallium (Fig. 1d, f), and hyperpallium (Fig. 1g, h, i). We quantified transgene expression by measuring the proportion of transgene-positive area in the injection site. For this, we extracted standardized 500 × 500 µm ROIs in areas with transgene expression for both AAV1-CAG-tdTomato and AAV8-CAG-tdTomato to compare expression levels.

**Figure 1.**
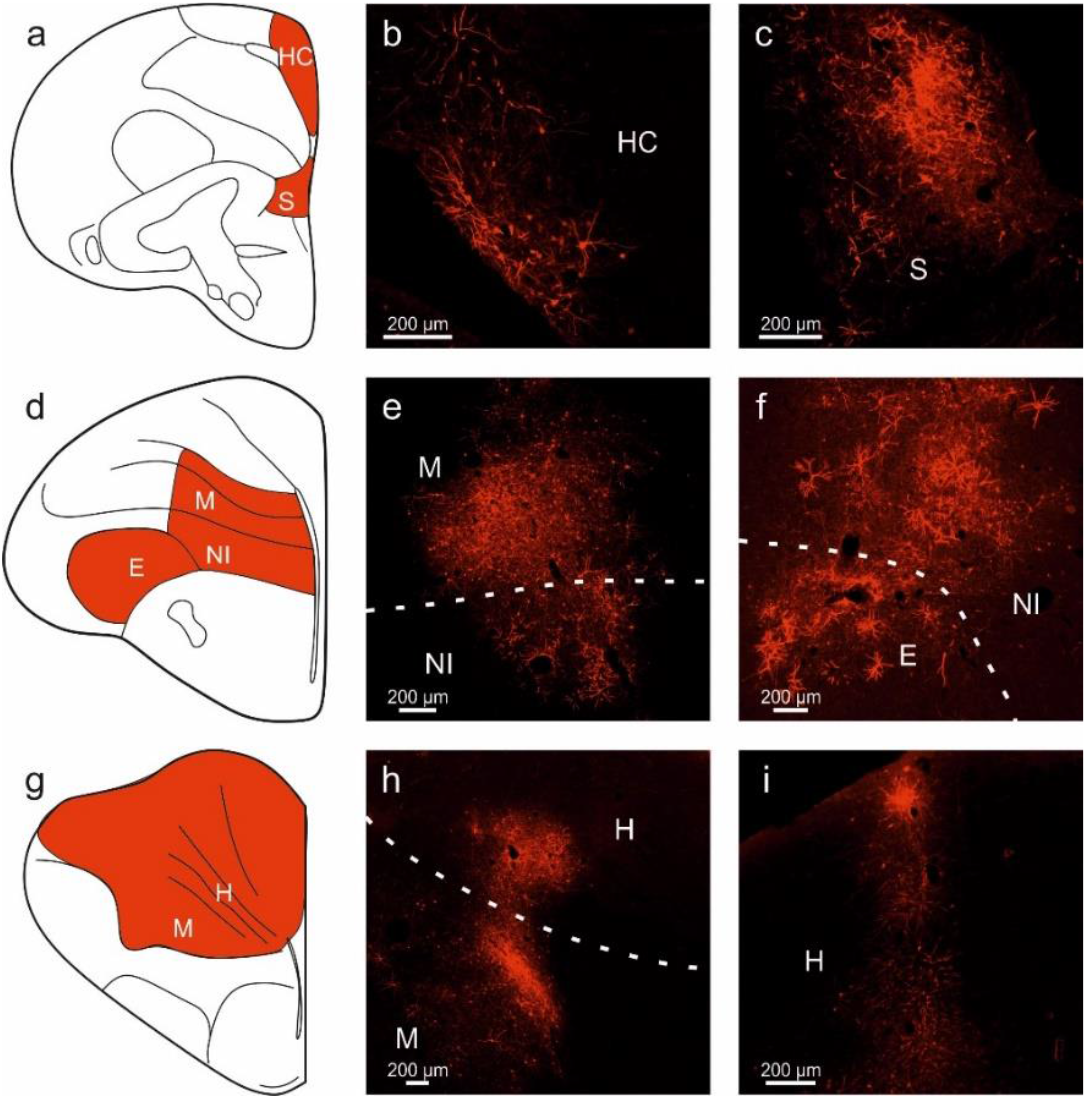
Transgene expression after AAV8 injection into various regions of the pigeon brain. (a, d, g) Schematic illustration of pigeon brain regions where expression was observable after injections of AAV8-CAG-tdTomato. Injections were performed in five hemispheres in three animals. Expression was found in the (b) hippocampus, (c) septum, (e, h) mesopallium, (e, f) nidopallium, (f) entopallium, and (h, i) hyperpallium. All scale bars represent 200 µm.

Using this method, we first quantified AAV1-CAG-tdTomato expression, which yielded a mean transduced area of 22.05% ± 1.65 SEM (n = 18 slices), closely matching previously reported values. AAV8-CAG-tdTomato resulted in a slightly lower mean transduced area of 15.43% ± 1.55 SEM (n = 14 slices).

Given previous reports of AAV8’s preference for non-neuronal cells [24,25], we performed double-labeling for RFP and NeuN, a neuronal marker, to determine the proportion of transduced neurons. Quantification of 22 brain slices from three animals, conducted independently by two examiners, revealed that 82.07% ± 1.39 SEM of transduced cells co-localized with NeuN, confirming they were neurons (Fig. 2a–d). To assess potential neurotoxicity, we examined GFAP expression. AAV8 did not induce GFAP expression at the injection sites (Fig. 2f–h). Notably, GFAP staining was visible in the hippocampus (Fig. 2e), a region where GFAP expression occurs naturally, confirming the staining’s validity. These findings indicate that AAV8 is both effective and non-toxic for neuronal gene transfer in pigeons.

**Figure 2.**
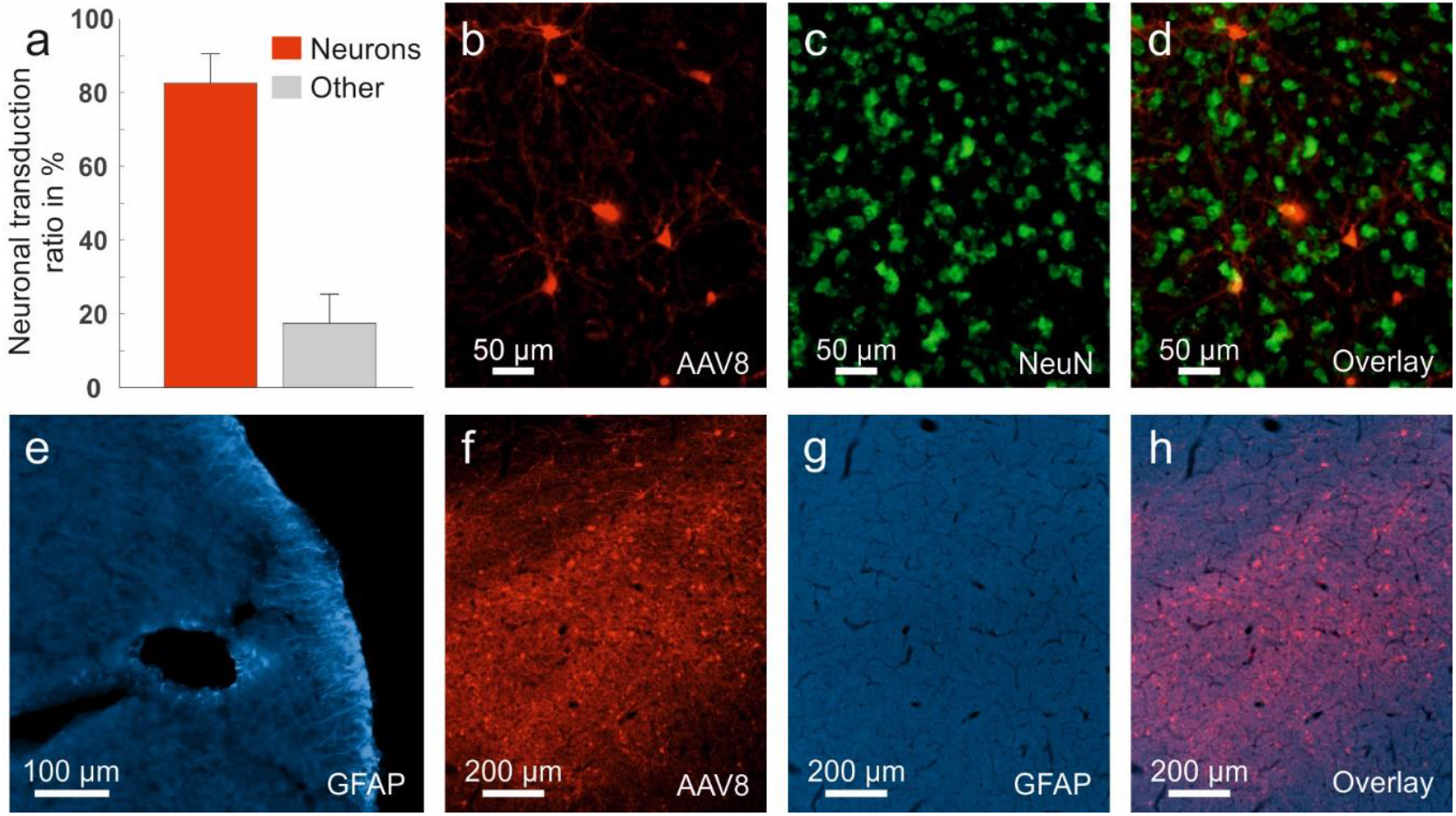
Neuronal specificity and neurotoxicity for AAV8. (a) Neuronal transduction ratio observed for injections with AAV8-CAG. (b-d) Fluorescence images depicting (b) RFP expression, (c) NeuN expression and (d) co-localization of RFP and NeuN. (e-h) Fluorescence images depicting (e) GFAP expression in a positive control area (the hippocampus), (f) RFP expression, (g) an absence of GFAP in the injection site and (h) co-localization of RFP and GFAP. Injections were performed in five hemispheres of three pigeons. The neuronal transduction ratio was determined in a total of 22 sections. Scale bars represent (b-d) 50 µm, (e) 100 µm or (f-h) 200 µm.

### Comparison of AAV Serotypes for Anterograde and Retrograde Transport

Anterograde and retrograde transport are crucial features for applications such as projection-specific optogenetics. Previous studies have shown that different AAV serotypes exhibit varying degrees of transport capability. For instance, AAV1 demonstrates robust anterograde and retrograde expression in pigeons [8]. Additionally, specialized variants like AAVretro have been engineered to enhance retrograde transport efficiency [26]. To evaluate these properties, we compared the performance of AAV8 (Fig. 3a–c), AAVretro (Fig. 3d–f), and AAV1 (Fig. 3g–i). Consistent with previous findings, AAV1 showed strong anterograde transgene expression, evidenced by fiber expression (Fig. 3h), and extensive retrograde labeling in cell bodies (Fig. 3i). AAV8 also demonstrated significant anterograde transgene expression, with labeled fibers observed in multiple areas (Fig. 3b), as well as retrograde labeling in the NCL following injections into the anterior mesopallium/nidopallium (Fig. 3c). In contrast, AAVretro exhibited minimal expression, with signal largely confined to the injection site (Fig. 3d–f). Retrograde labeling was virtually absent, with only sparse dendritic labeling detected in the nucleus rotundus following entopallium injections. These results suggest that AAVretro has limited utility in pigeons, while AAV8 and AAV1 are more effective for both anterograde and retrograde transport.

**Figure 3.**
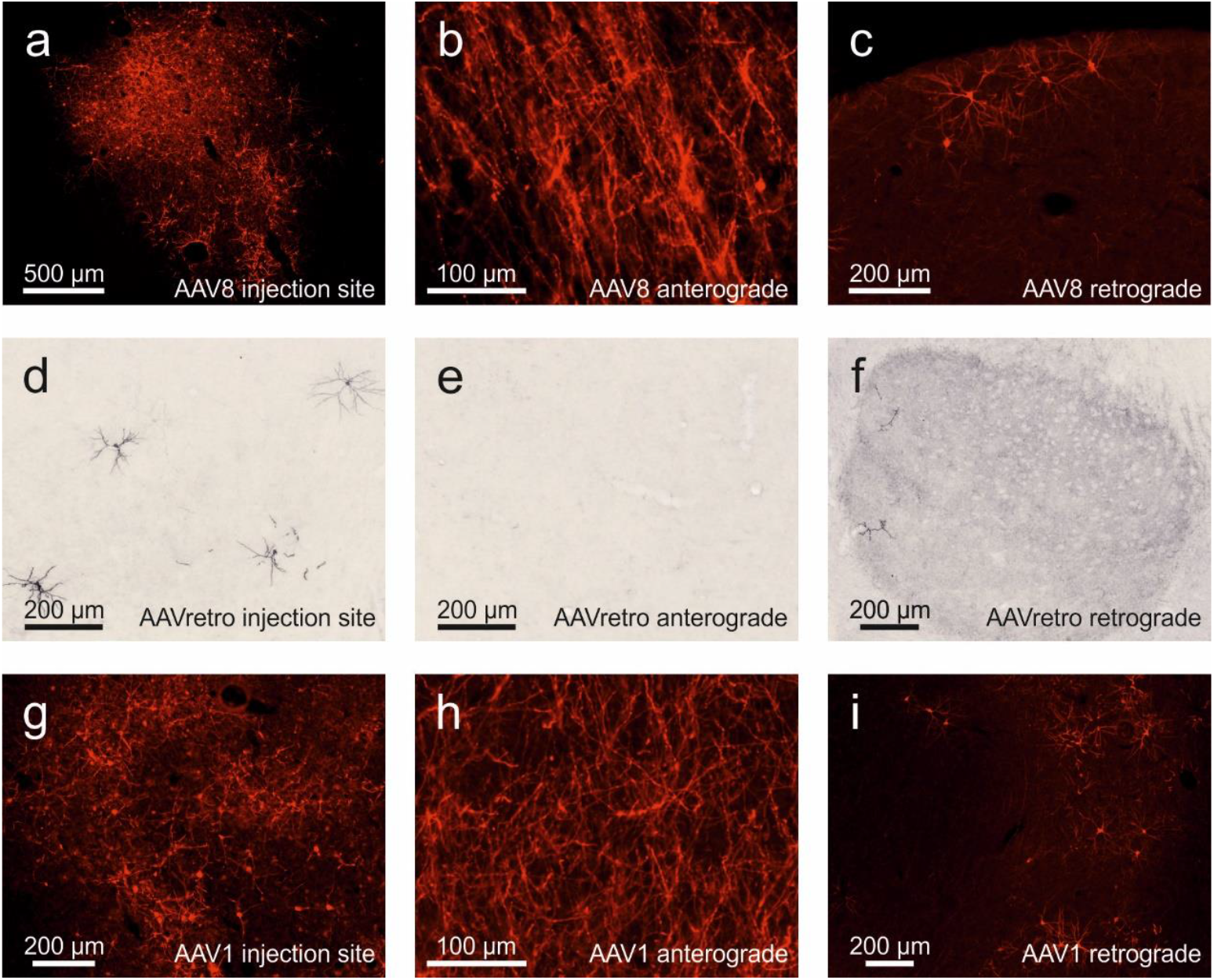
Anterograde and retrograde properties of AAV1, AAV8 and AAVretro. (a) Representative image of RFP expression at the injection site following AAV8 delivery, highlighting local transduction efficiency. (b) RFP-labeled fibers observed outside the injection site, indicating successful anterograde expression of the transgene. (c) RFP-positive cells identified in regions distal to the injection site, demonstrating retrograde transgene expression. (d) Representative image of ChR2 expression at the injection site following AAVretro delivery, highlighting local transduction efficiency. (e) No ChR2-labeled fibers were observed outside the injection site, indicating unsuccessful anterograde transgene expression. (f) Virtually no ChR2-positive cells could be identified in regions distal to the injection site such as the nucleus rotundus, demonstrating virtually no retrograde transgene expression. (g) Representative image of RFP expression at the injection site following AAV1 delivery, highlighting local transduction efficiency. (h) RFP-labeled fibers observed outside the injection site, indicating successful anterograde expression of the transgene. (i) RFP-positive cells identified in regions distal to the injection site, demonstrating retrograde transgene expression. The injected constructs were (a-c) AAV8-CAG-tdTomato (five hemispheres in three animals), (d-f) AAVretro (six hemispheres in three animals), and (g-i) AAV1-CAG-tdTomato (four hemispheres in two animals). Scale bars indicate (a) 500 µm, (b, h) 100 µm or (c-f, i) 200 µm.

### Validation of Cell-Type Specific Promoter/Enhancer Systems for Transgene Expression

We evaluated the specificity and usability of cell-type-specific promoter/enhancer systems for driving transgene expression in excitatory and inhibitory neurons. To this end, we paired AAV1 with either the CaMKIIα promoter, targeting CaMKIIα-positive excitatory neurons, or the mDLX enhancer, designed to drive expression in inhibitory interneurons [27,28]. To confirm specificity, brain sections from each vector were double-stained for CaMKIIα and GABA. For AAV1-CaMKIIα-NpHR, we observed strong transgene expression in several regions, including the entopallium, nidopallium, wulst, hippocampus, striatum and globus pallidus. Compared to the ubiquitous CAG promoter, expression levels were slightly lower, as expected for a cell-type-specific approach. Nevertheless, we found that 10.95 % ± 1.90 SEM CaMKIIα-positive neurons co-expressed the transgene. To determine the specificity of the CaMKIIα promoter, we determined the co-localization of transgene positive cells with either CaMKIIα or GABA. The co-localization analysis revealed a strong overlap with CaMKIIα-positive cells (84.57 % ± 1.87 SEM), with minimal overlap with GABA-positive cells (6.79 % ± 1.28 SEM) (Fig. 4a–f). While not entirely exclusive to CaMKIIα-positive neurons, the system exhibited a strong preference for this cell type. In contrast, for AAV1-mDLX-ChR2 we observed that 18.38 % ± 2.10 SEM of GABA-positive neurons were transgene-positive. Apart from that, transgene expression was predominantly localized to axons and dendrites in the entopallium, nidopallium and striatum. Co-localization analysis showed minimal overlap with CaMKIIα (8.22 % ± 3.78 SEM), while overlap with GABA-positive cells was higher (59.04 % ± 5.42 SEM) (Fig. 4g–l). These results suggest that while neither system achieves complete cell-type specificity, both exhibit clear preferences for their respective target cell populations.

**Figure 4.**
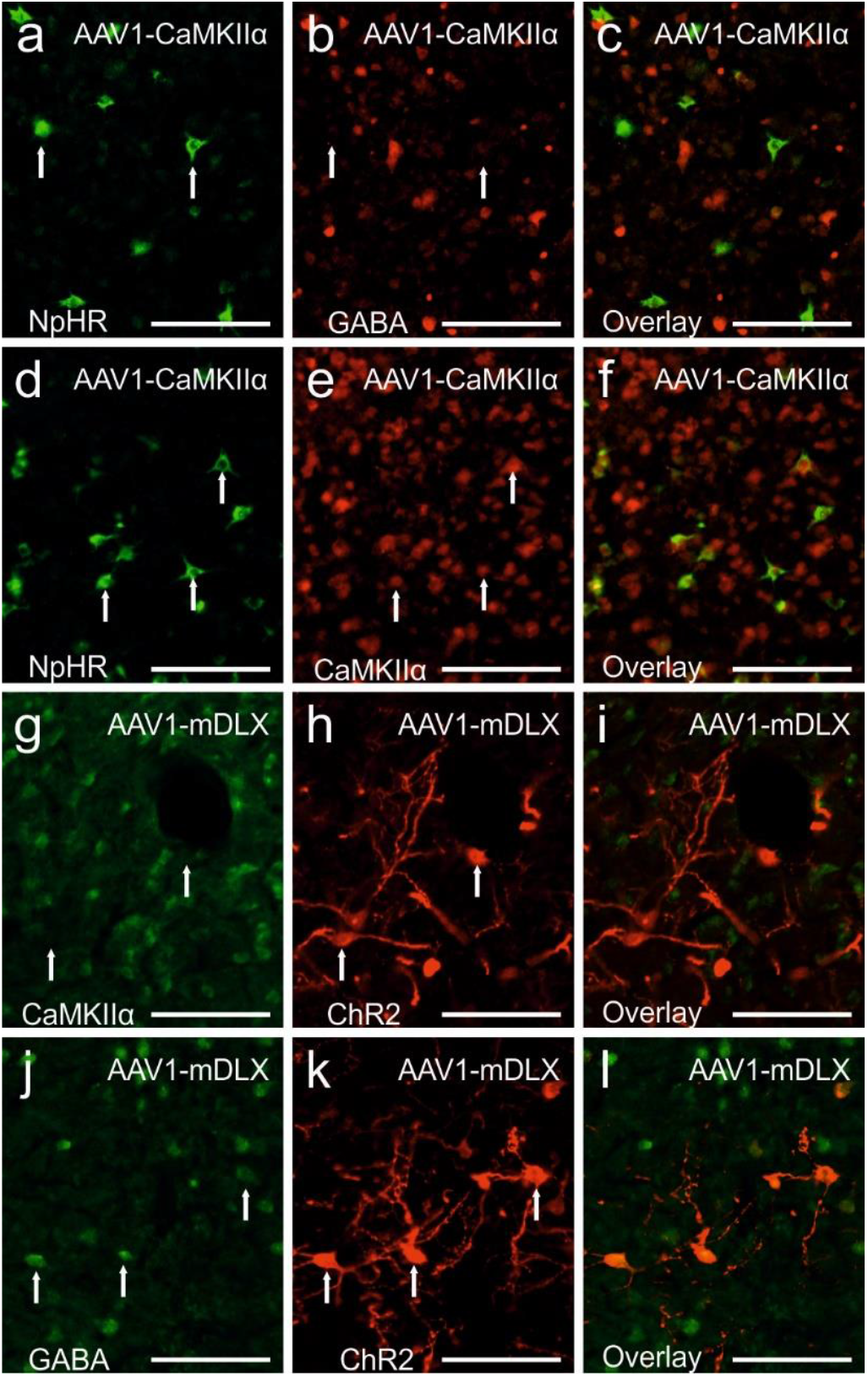
CaMKIIα and GABA specificity of CaMKIIα and mDLX promoter/enhancer systems. Fluorescence images of sections after injection of either (a-f) AAV1-CaMKIIα-eNpHR3.0-EYFP (n = eight hemispheres in five animals)or (g-l) AAV1-mDLX-ChR2-mCherry (n = six hemispheres in four animals). (a) Expression of NpHR after virus injection. (b) Fluorescence staining against GABA. (c) Overlay of NpHR and GABA signals showing no co-localizations. (d) Expression of NpHR after virus injection. (e) Fluorescence staining again CaMKIIα. (f) Overlay of NpHR and CaMKIIα showing co-localization of both signals. (g) Fluorescence staining against CaMKIIα. (h) Expression of ChR2 after virus injection. (i) Overlay of CaMKIIα and ChR2 signals showing no co-localization. (j) Fluorescence staining against GABA. (k) Expression of ChR2 after virus injection. (l) Overlay of ChR2 and GABA showing co-localization of both signals. Transgene expression was visualized with counterstainings against (a-f) YFP or (g-l) RFP. All scale bars represent 100 µm.

### Evaluation of Conditional and Advanced Expression Systems in Pigeons

Conditional expression using the Cre-loxP system offers precise control over transgene expression, making it particularly useful for optogenetic studies. In this approach, one vector expresses Cre recombinase, while a second vector contains a Cre-dependent promoter driving the transgene of interest. This ensures transgene expression only in the presence of Cre, enabling targeted manipulation of specific cells or projections [29]. To test this system, we injected AAV8-EF1a-dblf-mCherry alone or together with AAV8-CAG-Cre. In the absence of Cre, no mCherry expression was observed (Fig. 5a). However, co-injections with Cre resulted in high levels of mCherry expression (Fig. 5b,c,d,e), demonstrating the feasibility of this approach for optogenetic applications in pigeons. This system is particularly promising for projection-specific targeting.

**Figure 5.**
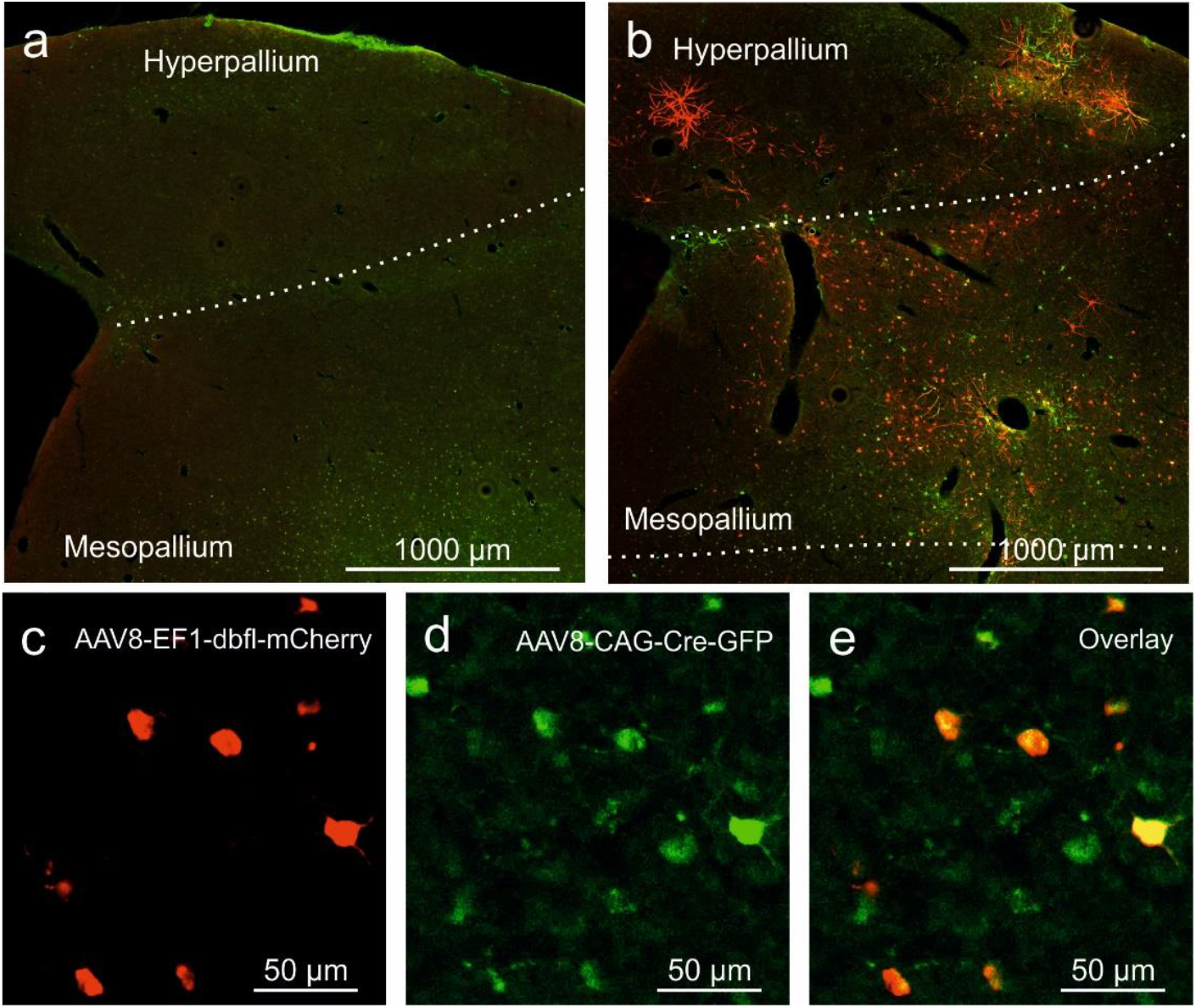
Conditional transgene expression using the Cre-LoxP system. (a) No transgene expression can be seen when AAV8-EF1-dbfl-mCherry is injected without the presence of Cre. (b) Simultaneous injection of AAV8-EF1-dbfl-mCherry and AAV8-CAG-Cre-GFP leads to expression of GFP (green) and mCherry (red). (c) Close-up image of mCherry expression. (d) Close-up image of GFP expression. (e) Co-localization of mCherry and GFP. Injections of AAV8-EF1-dbfl-mCherry were performed in four hemispheres in two animals, and AAV8-CAG-Cre-P2A-eGFP was injected in two hemispheres in two animals. Scale bars represent (a, b) 1000 µm or (c, d, e) 50 µm.

Additionally, we tested the Tet-Off system for inducible transgene expression in pigeons. The Tet-Off system, in which transgene expression is activated by default and downregulated by tetracycline, was implemented using AAV1-Syn1-tTa and three AAV1-TRE vectors encoding blue, yellow, and red fluorescent proteins (based on the TetBow technique; [30]. The TRE promoter is activated by the tetracycline-controlled transactivator (tTa) in the absence of tetracycline [31]. Following viral injections, transgene expression was achieved, with cells displaying mixed hues from multiple fluorescent proteins (Fig. 6). However, expression levels remained low after a six-week expression period, suggesting further optimization is needed for robust expression in pigeons.

**Figure 6.**
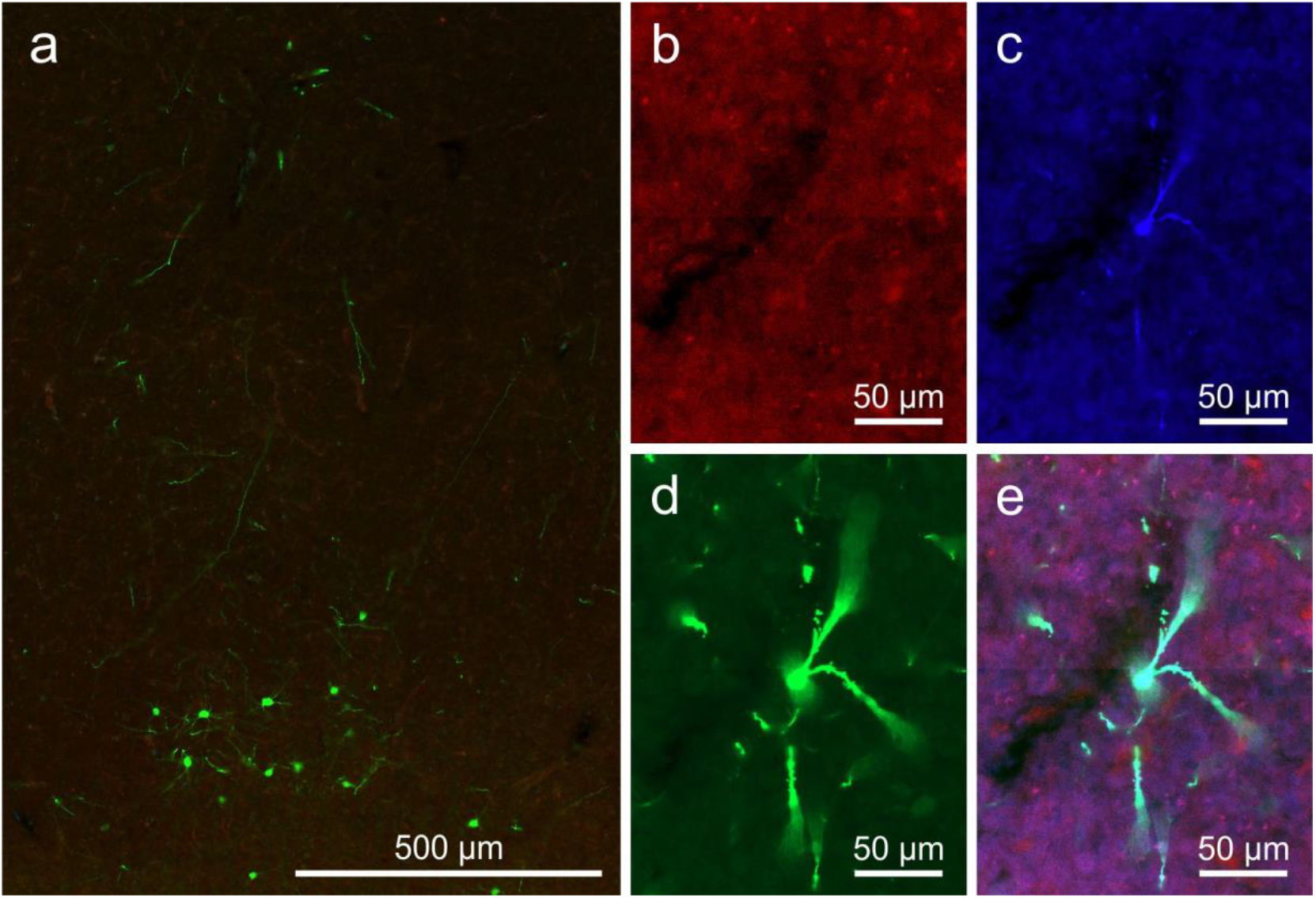
Tet-Off system can drive transgene expression in pigeons. (a) Transgene expression after viral injection. (b-e) Higher magnification images of a single cell expressing different levels of (b) tdTomato, (c) mTurquoise2, (d) eYFP, and (e) the overlay of b, c and d. Viral injections were performed in four hemispheres in two animals. Scale bars represent (a) 500 µm or (b-e) 50 µm.

## Discussion

This study set out to investigate the usability and effectiveness of various viral vectors to expand the available toolkit for transgene delivery in pigeons, a species for which transgenic approaches are not yet feasible. Specifically, we tested AAV8 and AAVretro for their ability to transduce cells, examined the specificity of the CaMKIIα promoter and mDLX enhancer, and provided proof-of-concept for conditional systems such as Cre-loxP and Tet-On/Tet-Off, laying the groundwork for future studies requiring precise gene transfer methods in this species.

### AAV8 expression characteristics

AAV8 effectively drove transgene expression across multiple brain regions in pigeons, including the nidopallium, mesopallium, hyperpallium, hippocampus, striatum, and septum, without inducing neurotoxic effects. In our study, GFAP staining for reactive gliosis and NeuN staining for neuronal integrity revealed no abnormalities at injection sites, indicating that AAV8 is well-tolerated in the pigeon brain. In contrast, previous studies reported neurotoxicity with AAV9 in pigeons [8], characterized by increased gliosis and decreased neuronal viability, highlighting species-specific differences in AAV serotype compatibility. These findings are consistent with rodent studies, where AAV8 is generally non-toxic when appropriately dosed but may induce neurotoxicity under certain conditions. For example, McLeod et al. (2024) found that AAV8 applications led to a mild inflammatory response, with a higher AAV titer correlating to more GFAP reactivity [33]. However, this inflammatory response did not result in a significant loss of neuronal tissue. Promoter choice also influences toxicity; ubiquitous strong promoters like CAG or CMV can exacerbate adverse effects, while cell-type-specific promoters may reduce toxicity by limiting transgene expression to targeted populations [34]. The absence of neurotoxicity in our study suggests that AAV8 offers a safer and more effective alternative to AAV9 for gene delivery in pigeons. This aligns with its favorable safety profile in rodents when optimized, reinforcing the need for species-specific testing to account for variations in vector performance and biocompatibility across models.

Moreover, we observed extensive anterograde transgene expression with labeling detected in regions distal to the injection sites, such as the arcopallium and midbrain structures. Retrograde transgene expression was also evident, although it appeared less pronounced than reported for AAV1 in earlier studies [8]. While some reports in rodents have noted that AAV8 exhibits weaker retrograde transgene expression compared to serotypes like AAV1 and AAV9 [35–37], the degree of retrograde transgene expression in our study is difficult to compare directly due to differences in target areas and experimental protocols. Our study used multiple injections, which may have obscured retrograde transport from specific sites. In contrast, AAVretro showed limited effectiveness in pigeons, with only sparse transgene expression observed in a small number of neurons and negligible retrograde labeling. While AAVretro was specifically engineered for retrograde transport and performs robustly in rodent models [26,37], it appears to lack utility in pigeons, potentially due to species-specific differences in neuronal architecture or viral tropism. On the other hand, AAV1 and AAV8 demonstrate strong anterograde and retrograde capabilities in pigeons, making them more suitable for experiments requiring retrograde labeling or distal transgene expression.

Interestingly, AAV8 has been reported to transduce non-neuronal cells, such as oligodendrocytes and astrocytes, more efficiently than other serotypes [24,25]. Our analysis showed that approximately 82% of transduced cells were NeuN-positive neurons, indicating strong neuronal tropism. This percentage is slightly lower than the 92% reported for AAV1 in pigeons [8], but still indicates a very strong tropism for neurons. This seems a little surprising given that AAV8 should be more efficient in also transducing non-neuronal cells. However, a similar neuronal tropism has been reported in other species for AAV8. For example, in marmosets, a study found up to 99% of cells transfected in the motor cortex, striatum or substantia nigra were neurons [38]. Interestingly, Klein et al. (2008) demonstrated that purification methods significantly impact AAV8 tropism [39]. Vectors purified with iodixanol showed a preference for neurons, while those purified with CsCl transduced astroglial cells more effectively. Our use of chloroform-purified vectors resulted in a primarily neuronal expression pattern, with a modest increase in transduction of non-neuronal cells compared to AAV1 (18% vs. 8%). These findings suggest that AAV8 may offer versatility for targeting neurons and non-neuronal cells in pigeons, depending on the desired application and the purification method employed.

### Cell type specificity

Another objective of this study was to evaluate viral vectors for cell-type-specific expression in pigeons, as previous work has been limited to the use of the broadly expressing CAG and CMV promoters and the neuron-specific hSyn promoter [8,23]. To expand the toolkit, we tested the specificity of the CaMKIIα promoter and the mDLX enhancer. The mDLX enhancer has demonstrated selectivity for GABAergic interneurons in multiple species, including mice, ferrets, gerbils, marmosets, and zebra finches [22,28]. This specificity likely arises from its derivation from the distalless homeobox 5 and 6 (Dlx5/6) genes, which are highly conserved across vertebrate species and are predominantly expressed in telencephalic interneurons [40,41].

We assessed the tropism of AAV1-mDLX in the pigeon brain through counterstaining for CaMKIIα and GABA. AAV1-mDLX exhibited overall weak expression levels and the co-localization analysis revealed a low overlap with CaMKIIα-positive neurons (8%) and a moderate overlap with GABA-immunoreactive cells (59%). This specificity aligns with expectations for mDLX but falls short of the over 90% co-localization with GABAergic neurons reported in other species [22]. A possible explanation for this discrepancy is the known difficulty in accurately detecting GABAergic neurons due to the rapid degradation of GABA and challenges with antibody binding during immunohistochemical staining [42]. Nevertheless, the minimal overlap with CaMKIIα-positive neurons, which far outnumber GABAergic neurons [28], supports the idea that mDLX maintains a preference for GABAergic neurons in pigeons.

However, the low overall expression levels achieved with AAV1-mDLX suggest that this enhancer may not be optimal for GABAergic-specific gene transfer in pigeons. A potential alternative is the glutamate decarboxylase 1 (GAD1) promoter, which was recently shown to drive GABAergic expression effectively in zebra finches. While the GAD1 promoter and mDLX enhancer produced comparable results in zebra finches, testing the GAD1 promoter in pigeons could provide a more reliable method for targeting GABAergic neurons [28].

We also evaluated the CaMKIIα promoter, which is widely used for targeting excitatory glutamatergic neurons. In our study, the CaMKIIα promoter drove abundant expression of NpHR across pigeon forebrain regions, showing a strong preference for CaMKIIα-positive cells. However, this specificity was not absolute, as some co-localization with GABAergic neurons was observed. While the CaMKIIα promoter has been considered selective for excitatory neurons, recent studies have raised concerns about its exclusivity [43–45]. Veres et al. (2023) [45] demonstrated that the CaMKIIα promoter could drive expression in specific subclasses of GABAergic neurons, even in cases where immunohistochemical staining failed to detect CaMKIIα. Moreover, functional evidence showed that these neurons could be activated when expressing ChR2, despite low fluorescent signal levels in histological analysis. This finding is surprising since GABAergic neurons typically lack detectable CaMKIIα expression [46,47]. One hypothesis is that the yCaMKII isoform, which is present in inhibitory neurons, may interact with the CaMKIIα promoter to facilitate gene expression [48].

Our results reflect this trend, as we observed some overlap with GABAergic neurons. However, the high specificity for CaMKIIα-positive neurons makes this promoter a viable option for targeting excitatory neurons compared to broad promoters like CAG. These findings underscore the importance of carefully validating the specificity of promoters for their intended applications in avian models.

### Conditional gene expression

The Cre-loxP system enables conditional and projection-specific gene expression [49]. This system requires two components: a viral vector encoding the Cre recombinase and another vector carrying a transgene flanked by loxP recognition sites. In our experiments, injections with only the double-floxed vector did not produce any detectable expression, confirming the necessity of Cre recombinase for activating the target gene. Injections that included both the Cre vector and the double-floxed vector led to abundant transgene expression, with clear colocalization of Cre and reporter proteins. While our results support the efficacy of the Cre-loxP system in pigeons, we acknowledge recent concerns about potential neurotoxic effects associated with AAV-mediated Cre expression. For example, Rezai Amin et al. (2019) observed significant cell death, reduced dopamine levels, and behavioral alterations following AAV-Cre injections into the substantia nigra of rodents [50]. In our study, NeuN and GFAP staining patterns did not reveal any signs of neurotoxicity, but we recommend researchers remain vigilant and conduct thorough histological assessments when employing this system.

The Tet-On/Tet-Off system offers another versatile method for conditional gene expression. By utilizing a tetracycline-controlled transactivator (tTa) and a tetracycline-responsive promoter (TRE), this system allows reversible control of transgene expression through the administration or withdrawal of tetracycline. We tested the Tetbow approach [30], which relies on the Tet-Off/tTa-TRE system to drive the expression of fluorescent proteins. While we successfully achieved transgene expression in the pigeon brain, the levels were substantially lower compared to other promoter systems after a waiting period of six weeks. Delayed expression could be attributed to species-specific differences in the time required for transgene activation. In pigeons, transgene expression generally appears slower than in rodents [8]. It is possible that the observed low fluorescence levels reflect an early stage of expression, as the TRE promoter requires sufficient tTa accumulation to initiate robust transcription. Extending the waiting period might improve expression levels. Beyond fluorescent labeling, the Tet-On/Tet-Off system provides exciting opportunities for reversible gene regulation. For instance, the addition of tetracycline to drinking water could allow researchers to toggle expression on and off, enabling precise temporal control over genetic modifications.

### Conclusions and Future Directions

This study provides key insights into the usability and limitations of viral vectors and genetic tools for transgene delivery in pigeons. AAV8 emerged as a robust candidate with strong neuronal tropism and anterograde/retrograde expression capabilities, while AAVretro proved ineffective. AAV1 and AAV8 remain reliable options for retrograde studies.

The CaMKIIα promoter demonstrated high specificity for excitatory neurons, making it suitable for targeting glutamatergic cells, though some expression in GABAergic neurons suggests caution in systems requiring complete exclusivity. In contrast, the mDLX enhancer showed limited expression, restricting its utility for behavioral studies. Future research should explore alternatives like the GAD1 promoter for improved targeting of inhibitory neurons.

Proof-of-concept demonstrations of Cre-loxP and Tet-On/Tet-Off systems confirm their potential for conditional and reversible gene expression in pigeons. However, delayed expression with the Tetbow approach highlights the need for species-specific optimization.

All in all, these tools expand the genetic toolkit for pigeons and lay the groundwork for advanced research in avian neuroscience.

## Methods

### Subjects

For this study 14 adult homing pigeons (*Columba livia*) were sourced from local breeders. The animals were individually housed, which food and water ad libitum, and subjected to a 12-hour light-dark cycle.

The experiments were carried out in compliance with the European Communities Council Directive of September, 22 2010 (2010/63/EU) and the specifications of the German law for the prevention of cruelty to animals and was approved by the animal ethics committee of the Landesamt für Natur, Umwelt und Verbraucherschutz (LANUV) NRW, Germany. We confirm that all methods were carried out in accordance with relevant guidelines and regulations and that the study was conducted in compliance with the ARRIVE guidelines.

### Viral vector production

Recombinant AAV stocks were generated using the triple-transfection technique and purified with chloroform [51,52]. Briefly, HEK293T cells were co-transfected with the vector plasmid, serotype plasmid, and a helper plasmid using polyethylenimine. Seventy-two hours post-transfection, the cells were collected by low-speed centrifugation, resuspended in lysis buffer (150 mM NaCl, 50 mM Tris-Cl, pH 8.5), and subjected to 5–7 freeze-thaw cycles. Following lysis, DNase I and MgCl_2_ were added, and the mixture was incubated at 37°C for 30 minutes. Polyethylene glycol (PEG-8000) was added to the supernatant to a final concentration of 10% (w/v), and the solution was incubated at 4°C for 2 hours. After centrifugation (3700 × g, 4°C, 20 minutes), the resulting PEG-precipitated pellet was resuspended in the clarified lysate.

For further purification, the resuspension was treated with additional PEG-8000, incubated at 4°C for at least 1 hour, and centrifuged again (3700 × g, 4°C, 20 minutes). The resulting pellet was dissolved in 50 mM HEPES buffer. Chloroform was then added in equal volume, the mixture was vortexed, and phase separation was achieved by centrifugation at room temperature (370 × g, 5 minutes). The aqueous layer was recovered, filtered through a 0.22 µm syringe filter, and further concentrated using PEG-8000. The final AAV preparation was resuspended in PBS containing 0.001% pluronic F68, aliquoted, and stored at −80°C.

### Viral vector injections

The injections of the viral vectors were applied locally to the brain regions of interest (the number of injections for each vector can be found in table 1). For the surgical procedures, anaesthesia was initiated with a combination of ketamine (Ketavet ®, Zoetis Deutschland GmbH, Germany, 60 mg/kg bodyweight IM), xylazine (Rompun ®, Bayer AG, Germany, 2 mg/kg body weight IM), and buprenorphine (Buprenovet ®, VetViva Richter GmbH, Austria, 0.5 mg/kg body weight IM) [53]. During the operation, gas anaesthesia using isoflurane (Forane 100%, Abbott GmbH & Co. KG, Wiesbaden, Germany) was applied additionally [53]. Before the procedure, the feathers on the head were cut to reveal the skin. Once the animals showed no pain reflexes and surgical tolerance was achieved, the birds were placed into the stereotactic apparatus. After an incision into the skin to expose the skull, craniotomies were drilled areas to reveal the brain tissue. The positions of the craniotomies depended on the location of the brain area. After removing the dura mater, the needle was inserted into the brain to perform the injections in the target regions. All injections were performed using a Hamilton syringe.

**Table 1:** Injected viral constructs.

| Construct | Manufacturer | Animals (n) | Hemispheres (n) | Injection volume | Expression time | Analysis | Staining method | Figure |
| --- | --- | --- | --- | --- | --- | --- | --- | --- |
| AAV8-CAG-tdTomato | Herlitze Lab | 3 | 5 | 5µl | 6 weeks | Neuronal tropism, anterograde/retrograde transport, general expression pattern | Fluorescence anti-RFP, anti-NeuN | 1, 2, 3 |
| AAV1-CAG-tdTomato | Addgene | 2 | 4 | 5µl | 6 weeks | Anterograde/retrograde transport | Fluorescence anti-RFP | 3 |
| AAVretro-CAG-ChR2 | Addgene | 3 | 6 | 5µl | 6 weeks | Anterograde/retrograde transport, general expression pattern | DAB anti-ChR2 | 3 |
| AAV1-CaMKIIα-eNpHR3.0-EYFP | Addgene | 5 | 8 | 5µl | 6 weeks | General expression pattern, GABA and CaMKIIα tropism | DAB anti-YFP; fluorescence anti-YFP, anti-GABA, anti-CaMKIIα | 4 |
| AAV1-mDLX-ChR2-mCherry | Addgene | 4 | 6 | 5µl | 6 weeks | General expression pattern, GABA and CaMKIIα tropism | Fluorescence anti-RFP, anti-ChR2, anti-GABA, anti-CaMKIIα | 4 |
| AAV8.CAG.Cre.P2A.eGFP.SV40.WPRE | Herlitze Lab | 2 | 2 | 5µl | 6 weeks | General expression pattern | Fluorescence anti-GFP | 5 |
| AAV8 EF1a.dbfl.mCherry.CW3SL | Herlitze Lab | 2 | 4 | 5µl | 6 weeks | General expression pattern, Cre-dependent expression | Fluorescence anti-RFP | 5 |
| AAV1-hSyn1-tTA | Addgene | 2 | 4 | 2.5µl | 6 weeks | General expression pattern | - | 6 |
| AAV1-TRE-mTurquoise2 | Addgene | 2 | 4 | 2.5µl | 6 weeks | General expression pattern | - | 6 |
| AAV1-TRE-EYFP | Addgene | 2 | 4 | 2.5µl | 6 weeks | General expression pattern | Fluorescence anti-YFP | 6 |
| AAV1-TRE-tdTomato | Addgene | 2 | 4 | 2.5µl | 6 weeks | General expression pattern | Fluorescence anti-RFP | 6 |

After 6 weeks to allow for transgene expression, the animals were transcardially perfused. Once a deep anaesthesia was achieved using equithesin (0.45ml per 100 g body weight), the perfusion was started by replacing the blood with a 0.9% sodium chloride (NaCl) solution, followed by fixation with 4° cold 4% paraformaldehyde (PFA) in 0.12M phosphate buffer (pH 7.4). Once the tissue was fixed, the brain was removed and stored in a postfix solution at 4° for 2 hours (4 % PFA and 30 % sucrose in phosphate buffered saline (PBS)). Afterwards the brains were stored in a 30 % sucrose solution for at least 24 hours to remove residual water for cryoprotection. For slicing, the brains were embedded in 15% gelatin/30% sucrose and again kept in postfix for 24 hours. The brains were sliced using a freezing microtome (Leica, Wetzlar, Germany) into 40 µm thick coronal sections.

### Immunohistochemistry

Every tenth slice of a brain series was used and all stainings were carried out with free-floating sections. As different staining combinations were used (see table 2), the following protocol describes the general staining procedure. In a first step, slices were rinsed in PBS for 2 × 5 min. Then, the slices were incubated in a 10 % blocking solution in PBS with 0.3% Triton-X-100 (PBST) for 30 min. Following the blocking step, the slices were transferred into a primary antibody solution (in PBST), where they remained for 72 h at 4 °C. After the primary incubation and further rinsing (3 × 10 min), the slices were then transferred into the solution containing the secondary antibodies (in PBST). This incubation step lasted for 1.5 h at room temperature. After a final rinsing step (3 × 10 min) the sections were mounted onto glass slides (Superfrost Plus, Thermo Scientific) and covered using Fluoromount-G (SouthernBiotech, Birmingham, USA).

**Table 2:** Antibody combinations used for different experiments.

| Experiment | Primary antibody | Manufacturer and RRID | Secondary antibody | Manufacturer and RRID |
| --- | --- | --- | --- | --- |
| **Assessment of AAV8 for Gene Transfer in Pigeon Brains** | rb anti-RFP 1:1000 | Antibodies-Online Cat# ABIN129578, RRID:AB_10781500 | gt anti-rabbit Alexa Fluor 594 1:500 | Thermo Fisher Scientific Cat# A32740, RRID:AB_2762824 |
|  | rat anti-GFAP 1:1000 | Innovative Research Cat#13-0300, RRID:AB_86543 | gt anti-rat Alexa Fluor 405 1:500 | Thermo Fisher Scientific Cat# A48261, RRID:AB_2890550 |
|  | ms anti-NeuN 1:500 | Thermo Fisher Scientific Cat# MA5-33103, RRID:AB_2802653 | gt anti-mouse Alexa Fluor 488 1:250 | Thermo Fisher Scientific Cat# A-11001, RRID:AB_2534069 |
| **Comparison of AAV Serotypes for Anterograde and Retrograde Transport** | ms anti-ChR2 1:1000 | Progen Cat#651180, RRID:AB_2892521 | biotinylated horse anti-mouse IgG 1:500 | Vector Laboratories Cat# BA-2000, RRID:AB_2313581 |
|  | rb anti-RFP 1:1000 | Antibodies-Online Cat# ABIN129578, RRID:AB_10781500 | gt anti-rabbit Alexa Fluor 594 1:500 | Thermo Fisher Scientific Cat# A32740, RRID:AB_2762824 |
| **Validation of Cell-Type Specific Promoter and Enhancer Systems for Transgene Expression** | ms anti-CaMKIIα 1:1000 | Abcam Cat# ab22609, RRID:AB_447192 | dk anti-mouse Alexa Fluor 488 1:500 | Thermo Fisher Scientific Cat# A-21202, RRID:AB_141607 |
|  |  |  | dk anti-mouse Alexa Fluor 594 1:500 | Thermo Fisher Scientific Cat# A-21203, RRID:AB_2535789 |
|  | rb anti-GABA 1:1000 | Thermo Fisher Scientific Cat# PA5-32241, RRID:AB_2549714 | dk anti-rabbit Alexa Fluor 488 1:500 | Thermo Fisher Scientific Cat# A-21206, RRID:AB_2535792 |
|  |  |  | dk anti-rabbit Alexa Fluor 594 1:500 | Thermo Fisher Scientific Cat# A-21207, RRID:AB_141637 |
|  | gt anti-YFP 1:1000 | SICGEN Cat# AB1166, RRID:AB_2895441 | dk anti-goat Alexa Fluor 488 1:500 | Thermo Fisher Scientific Cat# A-11055, RRID:AB_2534102 |
|  | rb anti-RFP 1:1000 | Antibodies-Online Cat# ABIN129578, RRID:AB_10781500 | dk anti-rabbit Alexa Fluor 594 1:500 | Thermo Fisher Scientific Cat# A-21207, RRID:AB_141637 |
|  | ms anti-ChR2 1:1000 | Progen Cat#651180, RRID:AB_2892521 | dk anti-mouse Alexa Fluor 594 1:500 | Thermo Fisher Scientific Cat# A-21203, RRID:AB_2535789 |
| **Evaluation of Conditional and Advanced Expression Systems in Pigeons** | rb anti-RFP 1:1000 | Antibodies-Online Cat# ABIN129578, RRID:AB_10781500 | dk anti-rabbit Alexa Fluor 594 1:500 | Thermo Fisher Scientific Cat# A-21207, RRID:AB_141637 |
|  | gt anti-YFP 1:1000 | SICGEN Cat# AB1166, RRID:AB_2895441 | dk anti-goat Alexa Fluor 488 1:500 | Thermo Fisher Scientific Cat# A-11055, RRID:AB_2534102 |

For DAB stainings, following the secondary antibody incubation and additional washing steps in PBS (3 × 10 min), the sections were then exposed to an avidin-biotin-peroxidase complex (1:100 in PBST, from the Vectastain Elite ABC kit) for 1 hour. Following additional PBS rinses (3 × 10 min), the sections were placed in a DAB working solution. This solution consisted of 5 mL of distilled water mixed with 4 drops (100 μL) of DAB stock, 2 drops (84 μL) of buffer stock, and 2 drops (80 μL) of nickel solution. The sections were then transferred into individual cell wells, each containing 1 mL of the prepared DAB solution. To initiate the reaction, 3 μL of H_2_O_2_ solution was added to each well. After 2 minutes of incubation, the sections were moved to wells containing PBS and rinsed twice for 5 minutes each. Finally, the sections were mounted onto gelatin-coated slides, dehydrated through a graded alcohol series, and coverslipped using Depex (Fluka, Munich, Germany). We verified antibody specificity based on either published literature or stainings in negative control sections. To verify reproducibility of our stainings, each staining was performed twice on separate brain series.

### Microscopic analysis

The stained slices were imaged using the ZEISS AxioImager M1 and ZEISS AxioScan.Z1 at 100x magnification with a Orca Flash 4.0 V3 camera. For the quantification of transduced cells and to verify co-localization of transduced cells with either NeuN, GABA or CaMKIIα, ZEN 3.5 lite was used to highlight transgene expressing cells and the respective co-localized proteins of interest.

For the quantification of transgene expression following AAV1 or AAV8 injections, as well as the proportion of transduced CaMKIIα or GABA cells, regions measuring 500×500µm were selected within the injection site of different regions. To analyze the transduced area for AAV1 and AAV8 as well as the proportion of transduced CaMKIIα-positive cells, we trained the automated segmentation software ZEN Intellesis (ZEN 3.7, Zeiss) with a 64 deep-feature model to recognize transgene expression or CaMKIIα. The trained Intellesis program was applied to segment all images, and quantification was performed using the ZEN 3.7 Image Analysis Wizard. We applied a filter for minimum area of 25 µm^2^ for the quantification of transgene expressing cells and of 10 µm^2^ for the quantification of CaMKIIα. For the analysis of the proportion of GABA-positive cells expressing the transgene following AAV1-mDLX injections, ZEN 3.5 lite was used to highlight GABA expressing cells and the co-localized transgene.

## Acknowledgements

This work was supported by the German Research Foundation (Deutsche Forschungsgemeinschaft, DFG) through grant SFB 1280 (A01 and A07) project number 316803389, the Federal Ministry of Education and Research (BMBF), the project AVIAN MIND, ERC-2020-ADG, LS5, GA No. 101021354, and the Max Planck Society.

## Author Contributions

N.R. conceived the experiments. T.S. and S.H produced viral vectors. K.H and J.M.T conducted viral injections and perfusions. N.R., J.M.T., K.H, and M.Z. performed the histology and microscopic analysis. O.G. and S.H acquired the funding. J.M.T wrote the original draft of the paper. N.R. edited the paper. All authors reviewed the paper.

## Data availability

The data will be made available from the corresponding author upon request.

## Competing interests statement

The authors declare no competing interests.

## Notes

### Competing Interest Statement

The authors have declared no competing interest.

## References

1. Balakhonov, D. & Rose, J. Crows Rival Monkeys in Cognitive Capacity. Scientific Reports 7, 8809 (2017).

2. Emery, N. J. & Clayton, N. S. The mentality of crows: convergent evolution of intelligence in corvids and apes. Science 306, 1903–1907 (2004).

3. Nieder, A. Evolution of cognitive and neural solutions enabling numerosity judgements: lessons from primates and corvids. Philosophical Transactions of the Royal Society B: Biological Sciences 373, 20160514 (2017).

4. Brainard, M. S. & Doupe, A. J. What songbirds teach us about learning. Nature 417, 351–358 (2002).

5. Jarvis, E. D. Learned birdsong and the neurobiology of human language. Annals of the New York Academy of Sciences 1016, 749–777 (2004).

6. Steinemer, A., Simon, A., Güntürkün, O. & Rook, N. Parallel executive pallio-motor loops in the pigeon brain. Journal of Comparative Neurology 532, e25611 (2024).

7. Biro, D., Freeman, R., Meade, J., Roberts, S. & Guilford, T. Pigeons combine compass and landmark guidance in familiar route navigation. Proceedings of the National Academy of Sciences of the United States of America 104, 7471–7476 (2007).

8. Rook, N. et al. AAV1 is the optimal viral vector for optogenetic experiments in pigeons (Columba livia). Commun Biol 4, 100 (2021).

9. Wallraff, H. G. Navigation by homing pigeons: updated perspective. Ethology Ecology & Evolution 13, 1–48 (2001).

10. Güntürkün, O. & Bugnyar, T. Cognition without Cortex. Trends in Cognitive Sciences 20, 291–303 (2016).

11. Stacho, M. et al. A cortex-like canonical circuit in the avian forebrain. Science 369 (2020).

12. Brenowitz, E. A. & Zakon, H. H. Emerging from the bottleneck: benefits of the comparative approach to modern neuroscience. Trends in Neurosciences 38, 273–278 (2015).

13. Li, X. et al. Fast noninvasive activation and inhibition of neural and network activity by vertebrate rhodopsin and green algae channelrhodopsin. Proceedings of the National Academy of Sciences of the United States of America 102, 17816–17821 (2005).

14. Aravanis, A. M. et al. An optical neural interface: in vivo control of rodent motor cortex with integrated fiberoptic and optogenetic technology. Journal of Neural Engineering 4, S143–56 (2007).

15. Galvan, A., Caiola, M. J. & Albaugh, D. L. Advances in optogenetic and chemogenetic methods to study brain circuits in non-human primates. Journal of Neural Transmission 125, 547–563 (2018).

16. Han, X. et al. Millisecond-timescale optical control of neural dynamics in the nonhuman primate brain. Neuron 62, 191–198 (2009).

17. Zhou, Z. C., Yu, C., Sellers, K. K. & Fröhlich, F. Dorso-Lateral Frontal Cortex of the Ferret Encodes Perceptual Difficulty during Visual Discrimination. Scientific Reports 6, 23568 (2016).

18. Roberts, T. F., Gobes, S. M. H., Murugan, M., Ölveczky, B. P. & Mooney, R. Motor circuits are required to encode a sensory model for imitative learning. Nature Neuroscience 15, 1454–1459 (2012).

19. Emiliani, V. et al. Optogenetics for light control of biological systems. Nature reviews. Methods primers 2 (2022).

20. Lisowski, L., Tay, S. S. & Alexander, I. E. Adeno-associated virus serotypes for gene therapeutics. Current Opinion in Pharmacology 24, 59–67 (2015).

21. Wu, Z., Asokan, A. & Samulski, R. J. Adeno-associated virus serotypes: vector toolkit for human gene therapy. Molecular Therapy 14, 316–327 (2006).

22. Dimidschstein, J. et al. A viral strategy for targeting and manipulating interneurons across vertebrate species. Nature Neuroscience 19, 1743–1749 (2016).

23. Nimpf, S., Kaplan, H. S., Nordmann, G. C., Cushion, T. & Keays, D. A. Long-term, high-resolution in vivo calcium imaging in pigeons. Cell reports methods 4, 100711 (2024).

24. Castle, M. J., Turunen, H. T., Vandenberghe, L. H. & Wolfe, J. H. Controlling AAV Tropism in the Nervous System with Natural and Engineered Capsids. Methods in molecular biology (Clifton, N.J.) 1382, 133–149 (2016).

25. Aschauer, D. F., Kreuz, S. & Rumpel, S. Analysis of transduction efficiency, tropism and axonal transport of AAV serotypes 1, 2, 5, 6, 8 and 9 in the mouse brain. PloS one 8, e76310 (2013).

26. Tervo, D. G. R. et al. A Designer AAV Variant Permits Efficient Retrograde Access to Projection Neurons. Neuron 92, 372–382 (2016).

27. Kaur, J. & Berg, R. W. Viral strategies for targeting spinal neuronal subtypes in adult wild-type rodents. Scientific Reports 12, 8627 (2022).

28. Spool, J. A. et al. Genetically identified neurons in avian auditory pallium mirror core principles of their mammalian counterparts. Current biology : CB 31, 2831-2843.e6 (2021).

29. Tye, K. M. & Deisseroth, K. Optogenetic investigation of neural circuits underlying brain disease in animal models. Nature Reviews Neuroscience 13, 251–266 (2012).

30. Sakaguchi, R., Leiwe, M. N. & Imai, T. Bright multicolor labeling of neuronal circuits with fluorescent proteins and chemical tags. eLife 7 (2018).

31. Gossen, M. & Bujard, H. Tight control of gene expression in mammalian cells by tetracycline-responsive promoters. Proceedings of the National Academy of Sciences of the United States of America 89, 5547–5551 (1992).

32. Zhang, Y. & Looger, L. L. Fast and sensitive GCaMP calcium indicators for neuronal imaging. The Journal of physiology 602, 1595–1604 (2024).

33. McLeod, F. et al. AAV8 vector induced gliosis following neuronal transgene expression. Frontiers in neuroscience 18, 1287228 (2024).

34. Xiong, W. et al. AAV cis-regulatory sequences are correlated with ocular toxicity. Proceedings of the National Academy of Sciences 116, 5785–5794 (2019).

35. Castle, M. J., Gershenson, Z. T., Giles, A. R., Holzbaur, E. L. F. & Wolfe, J. H. Adeno-associated virus serotypes 1, 8, and 9 share conserved mechanisms for anterograde and retrograde axonal transport. Human Gene Therapy 25, 705–720 (2014).

36. Cearley, C. N. & Wolfe, J. H. Transduction characteristics of adeno-associated virus vectors expressing cap serotypes 7, 8, 9, and Rh10 in the mouse brain. Molecular Therapy 13, 528–537 (2006).

37. Wang, J. & Zhang, L. Retrograde axonal transport property of adeno-associated virus and its possible application in future. Microbes and infection 23, 104829 (2021).

38. Masamizu, Y. et al. Local and retrograde gene transfer into primate neuronal pathways via adeno-associated virus serotype 8 and 9. Neuroscience 193, 249–258 (2011).

39. Klein, R. L., Dayton, R. D., Tatom, J. B., Henderson, K. M. & Henning, P. P. AAV8, 9, Rh10, Rh43 vector gene transfer in the rat brain: effects of serotype, promoter and purification method. Molecular Therapy 16, 89–96 (2008).

40. Zerucha, T. et al. A highly conserved enhancer in the Dlx5/Dlx6 intergenic region is the site of cross-regulatory interactions between Dlx genes in the embryonic forebrain. The Journal of Neuroscience 20, 709–721 (2000).

41. Ghanem, N. et al. Regulatory roles of conserved intergenic domains in vertebrate Dlx bigene clusters. Genome Res. 13, 533–543 (2003).

42. Zhao, C., Eisinger, B. & Gammie, S. C. Characterization of GABAergic neurons in the mouse lateral septum: a double fluorescence in situ hybridization and immunohistochemical study using tyramide signal amplification. PloS one 8, e73750 (2013).

43. Keaveney, M. K. et al. CaMKIIα-Positive Interneurons Identified via a microRNA-Based Viral Gene Targeting Strategy. The Journal of Neuroscience 40, 9576–9588 (2020).

44. Watakabe, A. et al. Comparative analyses of adeno-associated viral vector serotypes 1, 2, 5, 8 and 9 in marmoset, mouse and macaque cerebral cortex. Neuroscience research 93, 144–157 (2015).

45. Veres, J. M., Andrasi, T., Nagy-Pal, P. & Hajos, N. CaMKIIα Promoter-Controlled Circuit Manipulations Target Both Pyramidal Cells and Inhibitory Interneurons in Cortical Networks. eNeuro 10 (2023).

46. Liu, X. B. & Jones, E. G. Localization of alpha type II calcium calmodulin-dependent protein kinase at glutamatergic but not gamma-aminobutyric acid (GABAergic) synapses in thalamus and cerebral cortex. Proceedings of the National Academy of Sciences of the United States of America 93, 7332– 7336 (1996).

47. Sík, A., Hájos, N., Gulácsi, A., Mody, I. & Freund, T. F. The absence of a major Ca2+ signaling pathway in GABAergic neurons of the hippocampus. Proceedings of the National Academy of Sciences of the United States of America 95, 3245–3250 (1998).

48. He, X. et al. Gating of hippocampal rhythms and memory by synaptic plasticity in inhibitory interneurons. Neuron 109, 1013-1028.e9 (2021).

49. McLellan, M. A., Rosenthal, N. A. & Pinto, A. R. Cre-loxP-Mediated Recombination: General Principles and Experimental Considerations. Current protocols in mouse biology 7, 1–12 (2017).

50. Rezai Amin, S. et al. Viral vector-mediated Cre recombinase expression in substantia nigra induces lesions of the nigrostriatal pathway associated with perturbations of dopamine-related behaviors and hallmarks of programmed cell death. Journal of neurochemistry 150, 330–340 (2019).

51. Grieger, J. C., Choi, V. W. & Samulski, R. J. Production and characterization of adeno-associated viral vectors. Nat Protoc 1, 1412–1428 (2006).

52. Eickelbeck, D. et al. CaMello-XR enables visualization and optogenetic control of Gq/11 signals and receptor trafficking in GPCR-specific domains. Commun Biol 2, 60 (2019).

53. Serir, A. et al. Balanced anesthesia in pigeons (Columba livia): a protocol that ensures stable vital parameters and feasibility during long surgeries in cognitive neuroscience. Frontiers in physiology 15, 1437890 (2024).

